# Mass cytometry analysis of the NK cell receptor-ligand repertoire reveals unique differences between dengue-infected children and adults

**DOI:** 10.1101/2020.07.27.223339

**Authors:** Julia L. McKechnie, Davis Beltrán, Anne-Maud M. Ferreira, Rosemary Vergara, Lisseth Saenz, Ofelina Vergara, Dora Estripeaut, Ana B. Araúz, Laura J. Simpson, Susan Holmes, Sandra López-Vergès, Catherine A. Blish

## Abstract

Dengue virus (DENV) is a significant cause of morbidity in many regions of the world, with children at the greatest risk of developing severe dengue. Natural killer (NK) cells, characterized by their ability to rapidly recognize and kill virally infected cells, are activated during acute DENV infection. However, their role in viral clearance versus pathogenesis has not been fully elucidated. Our goal was to profile the NK cell receptor-ligand repertoire to provide further insight into the function of NK cells during pediatric and adult DENV infection. We used mass cytometry (CyTOF) to phenotype isolated NK cells and peripheral blood mononuclear cells (PBMCs) from a cohort of DENV-infected children and adults. Using unsupervised clustering, we found that pediatric DENV infection leads to a decrease in total NK cell frequency with a reduction in the percentage of CD56^dim^CD38^bright^ NK cells and an increase in the percentage of CD56^dim^perforin^bright^ NK cells. No such changes were observed in adults. Next, we identified markers predictive of DENV infection using a differential state test. In adults, NK cell expression of activation markers, including CD69, perforin, and Fas-L, and myeloid cell expression of activating NK cell ligands, namely Fas, were predictive of infection. In contrast, NK cell expression of the maturation marker CD57 and increased myeloid cell expression of inhibitory ligands, such as HLA class I molecules, were predictive of pediatric DENV infection. These findings suggest that acute pediatric DENV infection may result in diminished NK cell activation, which could contribute to enhanced pathogenesis and disease severity.

## Introduction

Dengue virus (DENV), a flavivirus with four serotypes (DENV1-4), is the most prevalent arthropod-borne virus in the world. Infection begins when an individual is bitten by a DENV-infected *Aedes* mosquito. After an incubation period of four to ten days, a majority of DENV-infected individuals will develop an asymptomatic infection or mild symptoms associated with dengue fever such as fever, headache, vomiting, myalgia, and rash. Generally, these symptoms persist for three to seven days before patients enter into defervescence. However, upon defervescence a small percentage of patients develop severe dengue characterized by severe plasma leakage, hemorrhage, and/or organ impairment (1).

DENV infection presents differently in children and adults. Vomiting, skin rash, abdominal pain, and anorexia are commonly observed in children while myalgia, nausea, retro-orbital pain, arthralgia, headache, and leukopenia are symptoms typical of adult DENV infection (2–5). Interestingly, children under the age of 16 are not only more likely to develop symptomatic dengue; they are also more likely to develop severe dengue and succumb to the infection (2, 6–10). There are several potential reasons as to why this is the case. The increase in plasma leakage observed in DENV-infected infants and children could be explained by higher capillary fragility (11). Additionally, antibody-dependent enhancement (ADE) caused by waning maternal antibodies or secondary DENV infection may contribute to increased disease severity (2, 12–14). While increased capillary fragility and ADE could both be contributing factors, increased risk of severe dengue in children compared to adults may also be due to differences in the immune response.

The evolution of the immune system with aging, as well as its implications for antiviral immunity have been well studied (15, 16). Broadly, people are born with an immature immune system that, with age, matures and develops immunological memory to previously encountered viruses. Traditionally, immune experience is strictly thought to shape the B and T cell repertoire. However, a previous study has demonstrated that immune experience acquired throughout life results in an increase in the diversity of natural killer (NK) cells (17), an innate immune cell subset which acts as one of the first responders to viral infection. Furthermore, numerous studies in the past decade have also revealed the ability of NK cells to develop both antigen-dependent and antigen-independent immunological memory (18).

NK cells kill infected target cells via three mechanisms: degranulation with release of cytotoxic mediators, receptor-mediated apoptosis, and antibody-dependent cellular cytotoxicity (ADCC). NK cells are activated to kill or secrete cytokines based on activating and inhibitory signals received from germline-encoded receptors binding to their cognate ligands on potential target cells. While NK cells are known to be activated during DENV infection, particularly during the acute phase (19–23), it is unclear which NK cell subsets are actually responding. Some putative receptor-ligand interactions that may trigger an anti-DENV NK cell response, such as NKp44/E protein, KIR2DS2/NS3-HLA-C, and others have been reported (24–26). We and others have also shown that upregulation of HLA class I molecules by DENV-infected cells suppresses the NK cell response (27–29). Importantly, prior work investigating the role of NK cells during *in vivo* DENV infection has been limited to examining either pediatric or adult patients, but never both in parallel (20–23, 30).

Our goal was to determine whether NK cells in children and adults respond differently to acute DENV infection. Using a cohort of pediatric and adult DENV patients from Panama, a dengue-endemic country, we profiled the expression of NK cell receptors and their ligands by mass cytometry (CyTOF). We found that acute DENV infection in children leads to a decrease in NK cell frequency, shifts in the composition of the NK cell compartment, as well as NK cell maturation marked by increased CD57 expression. No changes in NK cell frequency occurred in adults. However, DENV infection did result in increased expression of NK cell activation and functional markers, CD69, perforin, and Fas-L. Finally, analysis of myeloid cell subsets identified by unsupervised clustering revealed that DENV infection leads to a more dramatic departure from baseline phenotype in children, with greater induction of ligands for inhibitory NK cell receptors than observed in adults. Overall, this work suggests that pediatric DENV infection may result in a more suppressed NK cell response compared to adult DENV infection.

## Materials and Methods

### DENV cohort and ethical statement

Adult and pediatric patients experiencing symptoms of acute DENV infection for five days or less were enrolled at public health institutions in Panama City and surrounding suburban areas in Panama from 2013 to 2015. Healthy control samples were collected from volunteers at Gorgas Memorial Institute of Health Studies (ICGES). Patients were considered positive for dengue infection if they had a positive RT-PCR result. Suspected DENV patients negative for DENV, CHIKV, ZIKV were considered undifferentiated febrile patients. All dengue patients were infected with DENV-2, with the exception of one adult patient who was infected with DENV-1. The Institutional Review Board (IRB) of the Hospital del Niño (CBIHN-M-0634) approved this study protocol. Committees of ICGES, CSS, Santo Tomas Hospital, and Stanford University confirmed the protocol.

### PBMC sample processing, storage, and thawing

PBMCs were collected using Ficoll-Paque. Following isolation, PBMCs were stored at −80°C for 24-72 hours in freezing media (90% FBS, 10% DMSO) before being transferred to liquid nitrogen. Prior to use, PBMCs were thawed in complete media (RPMI-1640, 10% FBS, 1% L-glutamine, 1% penicillin/streptomycin), centrifuged, and counted using a TC20™ automated cell counter (Bio-Rad). One million PBMCs from each donor were set aside and kept on ice for ligand staining. NK cells were isolated from the remaining PBMCs by negative selection using a human NK Cell Isolation Kit (Miltenyi). Following NK cell isolation, NK cells were centrifuged and counted. NK cell isolation was not performed if a donor had fewer than 2 million viable PBMCs.

### Mass cytometry staining, data acquisition, and processing

Cells were stained for mass cytometry as previously described (29). Briefly, in-house conjugated antibodies were made using Maxpar^⍰^ X8 Antibody Labeling Kits (Fluidigm), while pre-conjugated antibodies were purchased from Fluidigm and Thermo Fisher Scientific. PBMCs or isolated NK cells were stained using the viability marker cisplatin (Enzo Life Sciences) and barcoded using a two-of-four barcoding scheme. Barcoded samples were pooled and stained with surface antibody cocktails before fixation with 2% paraformaldehyde and permeabilization (eBioscience Permeabilization Buffer). Samples were then stained with an intracellular panel and incubated in iridium-191/193 intercalator (DVS Sciences) for up to a week. Before the analysis by mass cytometry, samples were washed and diluted in EQ Four Element Calibration Beads. FCS files were normalized, and calibration beads were removed using the ParkerICI Premessa package. Normalized data were then de-barcoded using the same Premessa package. FlowJo^⍰^ 10.2 was used to gate on cell subsets of interest. Samples with less than 50% viability by TC20™ automated cell counter were excluded from subsequent analysis.

### CyTOF data analysis

The open source statistical software R (https://www.r-project.org/, version 3.6.1) (45) was used to perform the CyTOF data analysis. To account for heteroskedasticity, the signal intensities were transformed using the hyperbolic sine transformation (cofactor equals to 5) prior to any downstream analysis.

#### Clustering

The package *CATALYST* (version 1.10.1) (46) was used to perform the unsupervised clustering. We used 14 lineage markers (refer to **Figure 1A** for details) for the PBMCs and 35 markers (refer to **Figure 2A** for details) for the isolated NK cells. The default parameters of the clustering function were used. Briefly, this clustering method combines two algorithms: *FlowSOM* (47) which clusters the data into 100 high-resolution clusters and *ConsensusClusterPlus* (48) which groups these high-resolution clusters into metaclusters. To determine the optimal number of metaclusters, we used the delta area plot provided by the package at the end of the clustering step (8 metaclusters for the PBMCs; 7 metaclusters for the isolated NK cells).

**Figure 1:**
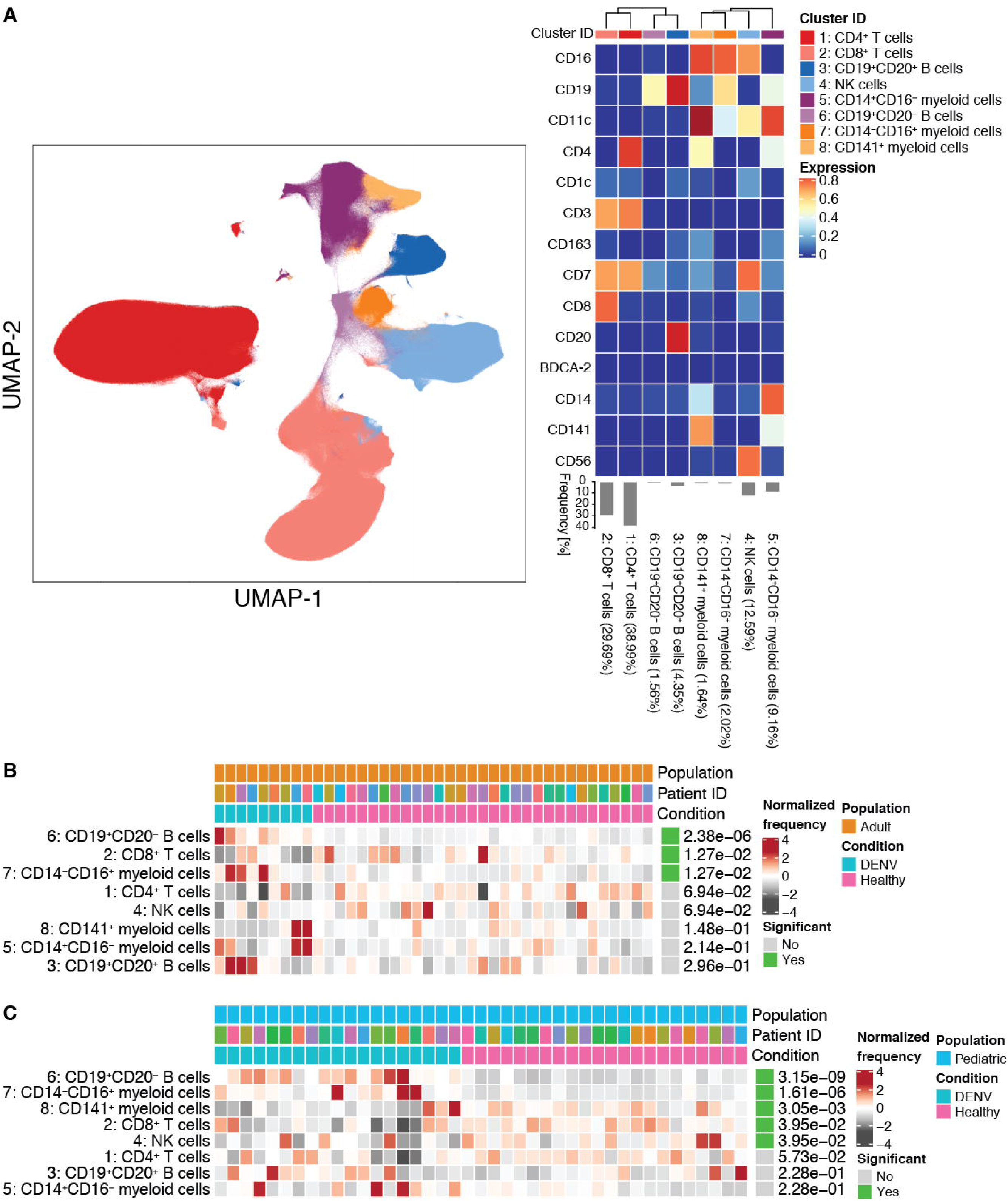
Acute DENV infection alters the frequencies of immune cell subsets. **(A)** UMAP visualization of PBMCs from DENV^+^, undifferentiated febrile illness, and healthy control pediatric and adult groups. The identity of each subset was determined based on expression of 14 lineage markers shown in the heat map. Each immune cell subset is color coded according to the key on the right. The heat map shows the expression of each marker (value scaled from 0 to 1) in each cluster. Clusters were hierarchically ordered based on similarity (dendrogram calculated using Euclidean distance as a metric and average as a linkage). The percentages associated with each cluster are the average of all pediatric and adult groups. **(B)** Results from differential abundance tests comparing the frequencies of immune cell subsets in DENV-infected adults (teal, n = 9) to healthy adults (pink, n = 31). **(C)** Results from a differential abundance test comparing the frequencies of immune cell subsets in DENV-infected children (teal, n = 19) to healthy children (pink, n = 22). Subsets whose frequencies were significantly different (adjusted p-value < 0.05) between the two states are denoted by green boxes. The proportions of each cluster in each sample are represented by the normalized frequencies. Grey boxes correspond to under-representation and red boxes correspond to over-representation. The frequencies were scaled using arcsine-square-root transformation and then z-score normalized in each cluster **(B and C)**.

**Figure 2:**
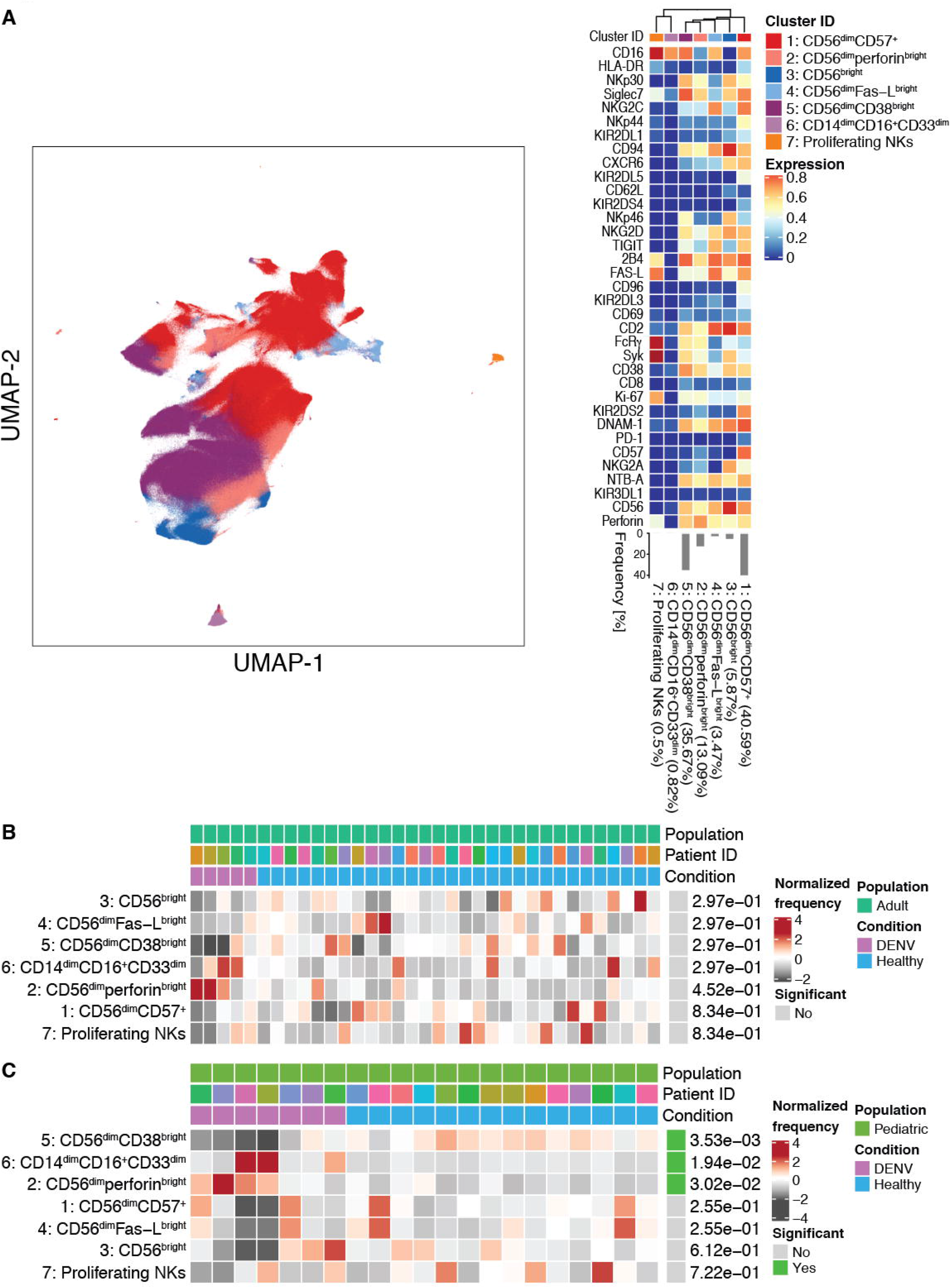
Acute DENV infection changes the composition of the NK cell compartment in children but not adults. **(A)** UMAP visualization of NK cells from DENV^+^ and healthy control pediatric and adult groups. The identity of each NK cell subset was determined based on the expression of 35 NK cell markers shown in the heat map. Each immune cell subset is color coded according to the key to the right of the heat map. The heat map shows the expression of each marker (value scaled from 0 to 1) in each cluster. Clusters were hierarchically ordered based on similarity (dendrogram calculated using Euclidean distance as a metric and average as a linkage). The percentages associated with each cluster are the average of all pediatric and adult groups. **(B)** Results from a differential abundance test comparing the frequencies of NK cell subsets in DENV-infected adults (purple, n = 5) to healthy adults (blue, n =30). **(C)** Results from a differential abundance test comparing the frequencies of immune cell subsets in DENV-infected children (purple, n = 7) to healthy children (blue, n = 14). NK cell subsets whose frequencies were significantly different (adjusted p-value < 0.05) between the two states are denoted by green boxes. The proportions of each cluster in each sample are represented by the normalized frequencies. Grey boxes correspond to under-representation and red boxes correspond to over-representation. The frequencies were scaled using arcsine-square-root transformation and then z-score normalized in each cluster **(B and C)**.

#### UMAP visualizations

We used the *uwot* package (version 0.1.5 available on CRAN) (49), which implements the Uniform Manifold Approximation and Projection (UMAP) algorithm (46). Using the same markers previously used for the clustering, we applied this dimensionality reduction method with the following parameters: 0.1 as the minimum distance; 20 as the number of nearest neighbors.

#### Differential abundance tests

Differential abundance tests were performed using the *diffcyt* package (version 1.6.1) (50) to identify differences in the frequencies of the cell clusters. We used the *diffcyt-DA-voom* function with the default parameters. Briefly, this method transforms the cluster cell counts to stabilize the mean/variance relationship with the *voom* method and then fits one Linear Model (LM) for each cluster. Each differential abundance test was performed on data which were filtered to the comparison of interest (adult or pediatric population; healthy or DENV-patients). The corresponding design matrix and contrast matrix were generated for each comparison. The reported p-values were adjusted by the False Discovery Rate (FDR) approach.

#### Differential state tests

To identify which markers were predictive of a specific state (for instance, healthy or DENV), we used the *CytoGLMM* package available on Github (51, 52). The method for unpaired samples implemented in this package utilizes a generalized linear model with bootstrap resampling to estimate the donor effect. The results of the model are the log-odds that a given marker is predictive of a specific state with a 95% confidence interval. The p-values are computed using Efron and Tibshirani 1993 methodology (53) and corrected for multiple testing by Benjamini-Hochberg method. 35 markers were considered for the differential state tests within the isolated NK cells (refer to **Figure 3** for details). 27 markers were considered for the differential state tests within the myeloid clusters of the PBMCs (refer to **Figure 4** for details). The following parameters were used: 2000 bootstraps, no subsampling was performed, and all samples were used (i.e. no threshold on the minimum number of cells per sample).

**Figure 3:**
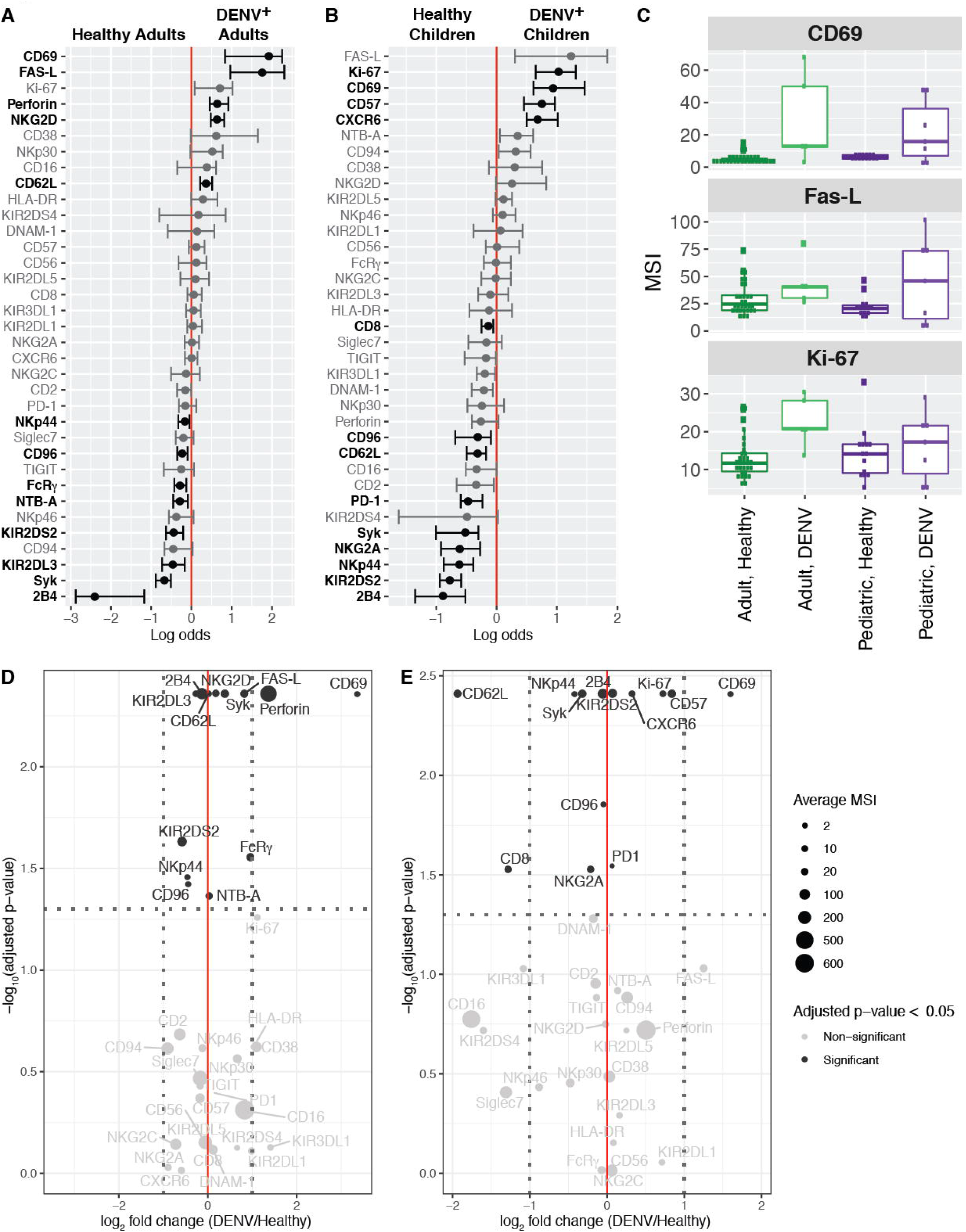
Acute DENV infection results in an activated NK cell phenotype in adults and NK cell maturation in children. **(A)** A generalized linear model with bootstrap resampling was used to identify markers on total NK cells that were predictive of adult DENV infection (right, n = 9) and healthy adults (left, n = 31). Black bars represent the 95% confidence interval. **(B)** A generalized linear model with bootstrap resampling was used to identify markers on total NK cells that were predictive of pediatric DENV infection (right, n = 19) and healthy children (left, n = 22). Markers with an adjusted p-value less than 0.05 are shown in black and markers with an adjusted p-values greater than 0.05 are shown in grey **(A and B)**. **(C)** Box plots showing the mean signal intensity (MSI) for the top three markers associated with DENV infection in both adults and children. Healthy adults (n = 30) are shown in dark green, DENV^+^ adults (n = 5) are shown in light green, healthy children (n = 14) are shown in dark purple, and DENV^+^ children (n = 7) are shown in light purple. **(D)** Volcano plot for adult NK cells illustrating markers whose adjusted p-values in **A** were less than 0.05 in black and markers whose adjusted p-values were greater than 0.05 in grey. The horizontal dashed line marks the 0.05 p-value cutoff. The −log_10_ p-value for each marker is shown on the y-axis. The DENV/healthy log_2_ fold change for each marker is shown on the x-axis. The two vertical dashed lines provide a reference point for markers whose expression is increased 2-fold. The size of each point corresponds to the mean signal intensity (MSI) for that specific marker. **(E)** Same as in **D**, but for pediatric NK cells. The MSIs are assigned according to the key at the right and they correspond to the raw (untransformed) MSI. The reported p-values are the adjusted p-values generated by the generalized linear model with bootstrap resampling **(D and E)**.

**Figure 4:**
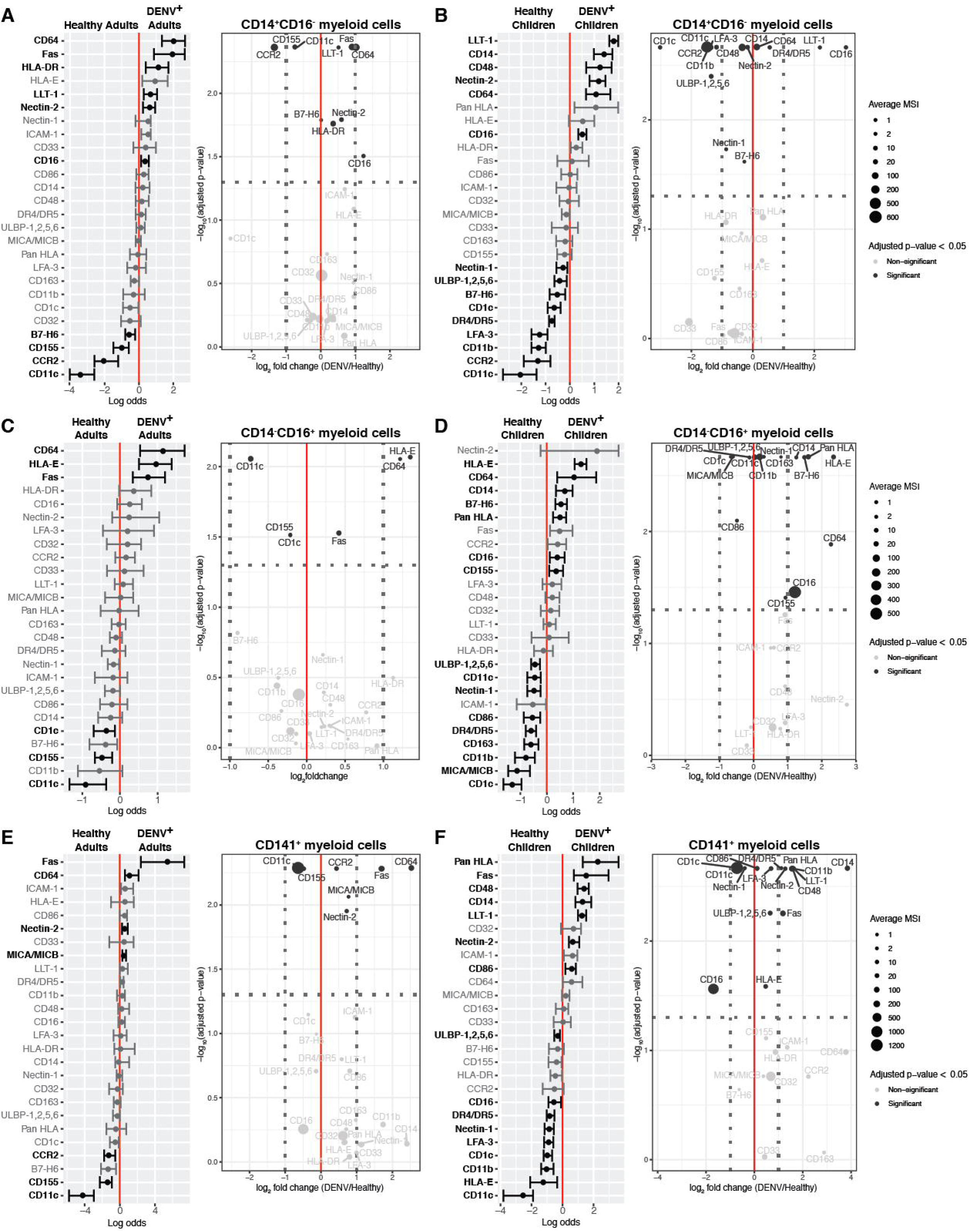
Pediatric DENV infection results in a dramatic shift in the phenotype of myeloid subsets and increased expression of inhibitory ligands compared to adult DENV infection. A generalized linear model with bootstrap resampling was used to identify markers on adult CD14^+^CD16^−^ **(A)**, CD14^−^CD16^+^ **(C)**, and CD141^+^ **(E)** myeloid cells and pediatric CD14^+^CD16^−^ **(B)**, CD14^−^CD16^+^ **(D)**, and CD141^+^ **(F)** myeloid cells that were predictive of DENV infection (right, n = 9 adults or 19 children) and healthy patients (left, n = 31 adults or 22 children). Volcano plots accompany each generalized linear model. The horizontal dashed line marks the 0.05 p-value cutoff. The −log_10_ p-value for each marker is shown on the y-axis. The DENV/healthy log2 fold change for each marker is shown on the x-axis. The two vertical dashed lines provide a reference point for markers whose expression is increased 2-fold. The size of each point corresponds to the mean signal intensity (MSI) for that specific marker. The MSI key is the same for adult and pediatric myeloid cells within the same subset and they correspond to the raw (untransformed) MSI. Both the generalized linear models and volcano plots illustrate markers whose adjusted p-values were less than 0.05 in black and markers whose adjusted p-values were greater than 0.05 in grey.

## Results

### Effects of DENV infection on immune cell subsets

Given the differences in disease course between DENV-infected children and adults, as well as the impacts of age on the antiviral immune response, we evaluated whether there were shifts in the abundances of immune cell subsets in Panamanian adult and pediatric DENV patients during the acute phase of infection. PBMCs were isolated from nine DENV^+^ adults, 31 healthy adults, 19 DENV^+^ children, and 22 healthy children **(Table I)**. All DENV cases were DENV2 cases, except for one adult DENV1 case. Samples were stained for CyTOF **(Supplementary Table I)** and gated down to live cells **(Supplementary Figure 1)**. Unsupervised clustering of live cells identified eight canonical immune cell subsets: CD4^+^ T cells (cluster 1, 38.99%), CD8^+^ T cells (cluster 2, 29.69%), CD19^+^CD20^+^ B cells (cluster 3, 4.35%), NK cells (cluster 4, 12.59%), CD14^+^CD16^−^ myeloid cells (cluster 5, 9.16%), CD19^+^CD20^−^ B cells (cluster 6, 1.56%), CD14^−^CD16^+^ myeloid cells (cluster 7, 2.02%), and CD141^+^ myeloid cells (cluster 8, 1.64%) **(Figure 1A)**. A differential abundance test revealed that DENV infection in both children and adults leads to a significant increase in the frequencies of CD19^+^CD20^−^ B cells and CD14^−^CD16^+^ myeloid cells as well as a decrease in CD8^+^ T cell frequency compared to healthy controls **(Figure 1B and 1C)**. Unlike adults, children also demonstrated decreases in the frequencies of CD141^+^ myeloid cells and NK cells, though there was heterogeneity between individuals **(Figure 1C)**. These data indicate that, in general, acute DENV infection has a broader impact on immune cell subsets in children than adults.

**Table I:**
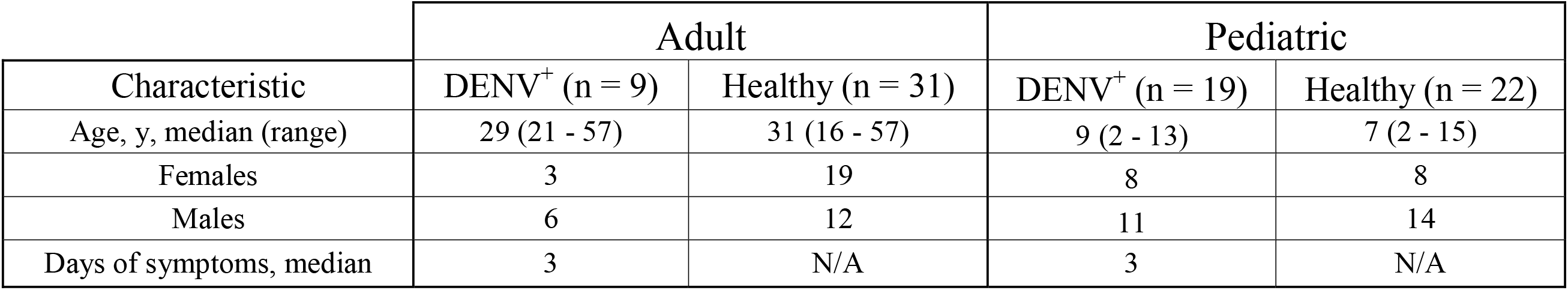
Panamanian DENV cohort demographics.

Our cohort also included seven children presenting with an undifferentiated febrile illness who were determined to be DENV^−^ by RT-PCR. To identify features associated with febrile illness, we performed a differential abundance test comparing these undifferentiated febrile patients to healthy children **(Supplementary Figure 2A)**. Similar to our DENV^+^ versus healthy children comparison, we observed significant increases in the frequencies of CD19^+^CD20^−^ B cells and CD14^−^CD16^+^ myeloid cells in undifferentiated febrile patients compared to healthy controls. An increase in CD14^+^CD16^−^ myeloid cell frequency and a decrease in CD4^+^ T cell frequency was also observed. Notably, unlike DENV infection, undifferentiated febrile illness had no impact on the frequency of NK cells when compared to healthy controls. To identify features associated explicitly with dengue virus infection, we performed a differential abundance test comparing DENV-infected children to undifferentiated febrile children. This revealed that the frequency of CD14^+^CD16^−^ myeloid cells was significantly lower in children infected with DENV **(Supplementary Figure 2B)**. Therefore, while both DENV infection and undifferentiated febrile illness uniquely impact certain immune cell subsets compared to healthy children, only a decrease in CD14^+^CD16^−^ myeloid cell frequency differentiates DENV infection from other febrile illnesses.

Given the importance of NK cells during the acute phase of viral infection, we were intrigued by the decrease in NK cell frequency observed in pediatric DENV cases. Consequently, we performed a deeper analysis of the NK cell compartment.

### Identification of NK cell subsets and the impact of DENV infection on their frequencies

To identify NK cell subsets responding to DENV infection, we performed unsupervised clustering of purified NK cells. NK cells were isolated from whole PBMCs and stained with an NK cell-focused CyTOF panel **(Supplementary Table II)**. Negative gating was performed before downstream analysis to remove any residual, non-NK cells **(Supplementary Figure 3)**. Our analysis identified the canonical CD56^bright^ and CD56^dim^ NK cell subsets **(Figure 2A)**. While CD56^bright^ NK cells clustered together (cluster 3, 5.87%), CD56^dim^ NK cells were divided amongst four clusters. The largest CD56^dim^ clusters (clusters 1 and 5) consisted of CD56^dim^CD57^+^ (40.59%) and CD56^dim^CD38^bright^ (35.67%) NK cells, respectively. The less abundant CD56^dim^ subsets corresponding to clusters 2 (13.09%) and 4 (3.47%) consisted of CD56^dim^perforin^bright^ and CD56^dim^Fas-L^bright^ NK cells, respectively. Two clusters, clusters 6 and 7, each made up less than 1% of the total NK cells. Of the 35 NK cell markers examined, cluster 6 only expressed one: CD16, making it likely that this cluster represents CD14^dim^CD16^+^CD33^dim^ monocytes that were not removed by purification or negative gating **(Supplementary Figure 3)**. Cells in cluster 7 express the NK cell markers 2B4, Fas-L, FcR□, Syk, Ki-67, and CD16. Therefore, it is likely that this cluster represents a small population of actively proliferating NK cells.

Next, we performed a differential abundance test to determine whether the frequencies of the identified NK cell subsets change during acute DENV infection. Interestingly, there were no significant differences in the composition of the NK cell compartment between DENV-infected adults and healthy adult controls **(Figure 2B)**. However, in children, DENV infection resulted in a significant decrease in the frequency of CD56^dim^CD38^bright^ NK cells and an increase in the frequency of CD56^dim^perforin^bright^ NK cells **(Figure 2C)**. There was also an increase in the abundance of the second smallest cluster (cluster 6), which, as previously discussed, is likely made up of CD14^dim^CD16^+^CD33^dim^ monocytes.

### Effects of DENV infection on NK cell phenotype

After observing significant shifts in the pediatric NK cell compartment upon DENV infection, we performed a differential state test to compare the phenotype of total NK cells in DENV-infected adults and children to their respective healthy controls **(Figure 3A and 3B)**. Interestingly, three markers of NK cell activation, CD69, Fas-L, and Ki-67, were associated with acute dengue in both adult and pediatric patients. These results are consistent with what has been previously published (19, 20, 22, 23). However, the extent to which NK cells upregulate these proteins varies between adults and children **(Figure 3C)**. There is also some heterogeneity within each of the DENV-infected groups.

We then visualized our NK cell CyTOF data using volcano plots, allowing us to compare each marker’s mean signal intensity (MSI), fold change (DENV/healthy), and how predictive it was of either state (DENV^+^ or healthy). As expected, DENV-infected adults had a greater than 10-fold increase in CD69 expression **(Figure 3D)**. Adult DENV infection was also characterized by robust expression of perforin, as well as a modest increase in expression of death ligand Fas-L and activating receptor NKG2D. CD69, perforin, Fas-L, and NKG2D were all predictive of adult DENV-infection. These data suggest that adult NK cells are activated upon DENV infection and likely capable of responding to DENV-infected cells.

Interestingly, DENV infection in children seemed to result in a mature NK cell phenotype **(Figure 3E)**. Expression of the maturation marker CD57 increased nearly 2-fold in DENV-infected individuals and was significantly predictive of infection, while expression of the immaturity marker NKG2A was reduced. CD69 was also upregulated in DENV-infected children, albeit to a lesser extent than in DENV-infected adults (2.8-fold vs 10-fold). Other proteins whose expression increased upon DENV infection in children were chemokine receptor CXCR6 and proliferation marker Ki-67. The expression of death receptor, PD-1, was also slightly increased. CD57, CD69, CXCR6, Ki-67, and PD-1 were all significantly predictive of pediatric DENV infection. Together, these data suggest that DENV infection may drive maturation and modest activation of NK cells in children.

### Myeloid cell expression of ligands for NK cell receptors

Given our finding that NK cells are activated to some extent during acute DENV infection irrespective of patient age, and our identification of a CD56^dim^perforin^high^ NK cell subset, we next sought to identify ligands for NK cell receptors that could contribute to their activation. As myeloid cells are the main targets of DENV infection (31), we focused on the three myeloid subsets identified by our unsupervised clustering: CD14^+^CD16^−^, CD14^−^CD16^+^, and CD141^+^ myeloid cells. In the adult CD14^+^CD16^−^ subset there were ten markers whose expression was significantly predictive of either DENV infection or healthy controls **(Figure 4A)**. The markers that were upregulated during DENV infection included death receptor Fas, inhibitory CD161 ligand LLT-1, and HLA-DR. In contrast, there were 15 markers whose expression was significantly altered in children **(Figure 4B)**. Markers that were upregulated during pediatric DENV infection included LLT-1 and death receptor 4/5 (DR4/DR5).

In the CD14^−^CD16^+^ subset there were six markers whose expression was significantly altered in adults **(Figure 4C)**. The markers whose expression was upregulated during DENV infection included Fas and HLA-E, whose role in suppressing the NK cell response to DENV-infected cells is currently unclear (29). In the pediatric CD14^−^CD16^+^ myeloid cell subset there were 16 significant markers **(Figure 4D)**. Those whose expression was higher during DENV infection included the well-known NK cell inhibitory ligands, HLA class I molecules, as well as known activating NKp30 and NKG2D ligands, B7-H6 and ULBP-1,2,5,6 respectively.

Finally, in the CD141^+^ myeloid subset there were seven significant markers in adults **(Figure 4E)**. Those that were upregulated during DENV infection included Fas as well as MICA/MICB, ligands for the activating NK cell receptor NKG2D. There were 16 significant markers in children **(Figure 4F)**. Inhibitory ligands DR4/DR5, LLT-1, and HLA class I molecules as well as activating ligands Fas and ULBP-1,2,5,6 along with the costimulatory marker CD86 were among those upregulated during pediatric DENV infection. These findings suggest that pediatric DENV infection leads to more significant changes in myeloid cell phenotype compared to adult infection. The phenotype of adult myeloid cells likely promotes NK cell activation, while the phenotype of pediatric myeloid cells likely dampens NK cell activation.

Based on the work done by Costa et al. demonstrating the importance of DNAM-1, 2B4, LFA-1, and CD2 in mediating the NK cell response to DENV-infected cells, we also looked at the impact of DENV infection on expression of their respective ligands (26). We found no consistent pattern in DNAM-1 ligands, Nectin-2 and CD155 (PVR), being associated with either state: DENV^+^ or healthy. The same was true for 2B4 and CD2 ligands, CD48 and LFA-3 respectively. Expression of ICAM-1, the LFA-1 ligand, was never significantly predictive of either state.

## Discussion

While severe dengue has historically been considered a children’s disease, there has been a recent increase in the average age of reported dengue cases, particularly in Southeast Asia (7, 32, 33). Considering that the geographical range of DENV is expected to expand in the coming years (34), increasing the portion of the population at risk of infection, it is important to determine whether there are age-related differences in how the immune system responds to DENV infection. Previous studies investigating the immune response to DENV infection have reported an increase in activated NK cells (19, 20, 22, 23, 30), suggesting a role for this immune cell subset. However, all of these studies solely focused on either pediatric or adult DENV infection, not both. Here we evaluated the NK cell receptor-ligand repertoire of pediatric and adult DENV patients by CyTOF. We show that, compared to adults, children have a lower frequency of NK cells and significant changes in the composition of their NK cell compartment. While DENV infection leads to a markedly activated NK cell phenotype in adults, it results in NK cell maturation and modest activation in children. Additionally, acute pediatric DENV infection leads to significant myeloid cell upregulation of inhibitory NK cell ligands, which is largely absent in adult DENV patients. Together, these findings suggest that compared to adults, DENV infection in children favors a more suppressed NK cell response.

Increases in monocyte, activated T cell, NK cell, and plasmablast frequencies as well as decreases in the frequencies of total CD3^+^ and CD4^+^ T cells during acute DENV have been previously reported (30, 35, 36). We observed similar results in our cohort with DENV-infected children and adults experiencing a decrease in the frequency of CD8^+^ T cells and increases in the frequencies of CD19^+^CD20^−^ B cells and CD14^−^CD16^+^ myeloid cells. In addition, we observed a decrease in the frequencies of CD141^+^ myeloid cells and NK cells during acute pediatric DENV infection. These changes in the CD141^+^ myeloid cell and NK cell compartments were not observed when children presenting with an undifferentiated febrile illness were compared to healthy children. One subset that is notably absent from our clustering analysis is CD14^+^CD16^+^ myeloid cells. Studies have found that the frequency of CD14^+^CD16^+^ monocytes increases during DENV infection and may stimulate differentiation of plasmablasts (37, 38). Our inability to detect this subset could be due to its low frequency compared to the other two monocyte subsets (39) or the fact that expansion of CD14^+^CD16^+^ monocytes typically occurs within the first two days of infection before dramatically dropping off (38).

Unsupervised clustering of isolated NK cells identified six NK cell subsets. Of the NK cell subsets identified, one expressed high levels of the functional marker perforin. In addition to perforin, the CD56^dim^perforin^bright^ subset expressed NKp30, Siglec-7, CD94, 2B4, CD2, FcR□, CD38, Ki-67, DNAM-1, and NTB-A. CD38 and Ki-67 expression suggests that this is an activated NK cell subset. This is further supported by the fact that these cells also express Siglec-7, which has been reported as a marker of highly functional NK cells (40). Moreover, *in vitro* studies have identified 2B4, CD2, and DNAM-1 as playing significant roles in mediating NK cell interactions with DENV-infected monocyte-derived dendritic cells (mDCs) (26). Blocking these proteins with antibodies resulted in decreased NK cell expression of CD69 as well as an increase in viral replication. This suggests that CD56^dim^perforin^bright^ NK cells are activated and may be capable of killing DENV-infected cells. NKp30 and DNAM-1 potentially play a role in mediating this activation. While the frequencies of the individual NK cell subsets are unchanged during adult DENV infection, the frequency of CD56^dim^perforin^bright^ NK cells increases in DENV-infected children suggesting a shift towards a degranulation response. Degranulation is an effective mechanism for killing virally infected cells. However, it is possible that in children this response is not specific enough, contributing to pathogenesis rather than viral clearance.

Using adult cohorts, Azeredo et al. and Gandini et al. have shown an association between NK cell activation and mild dengue (20, 41). Conversely, Green et al. observed a higher frequency of CD69^+^ NK cells in children who developed dengue hemorrhagic fever (DHF) compared to children with mild disease (19). While our cohort does not contain any patients classified as having DHF, our data supports this idea of NK cells responding differently to DENV infection in children versus adults. We found that expression of NK cell activation and functional markers NKG2D, Syk, Fas-L, perforin, and CD69 was higher in adult dengue patients compared to healthy controls and that expression of all these makers was predictive of adult DENV infection. While CD69 expression was increased in DENV-infected children and predictive of infection, the NK cell phenotype was also characterized by CD57 expression. Importantly, NKG2A expression was higher in healthy children and significantly predictive of that state. CD57 expression is induced upon NK cell stimulation and increases with age while NKG2A expression decreases with age (17, 42, 43). CD57 also defines a subset of NK cells that have a greater cytotoxic capacity and higher sensitivity to CD16 signaling (44). These data suggest that DENV infection induces NK cell maturation in children. In all likelihood, the adults in our cohort have greater immune experience than the children. Consequently, their NK cells are already mature and can mount a more rapid, functional response that may limit viral spread more effectively.

Importantly, our unsupervised clustering analysis complements the differential state tests we performed using a generalized linear model. Occasionally the results from these different analyses appear contradictory. For example, perforin expression was upregulated during adult DENV infection, however there was no corresponding increase in the frequency of the CD56^dim^perforin^bright^ NK cell subset. Similarly, expression of CD57 was upregulated during pediatric DENV infection, however no increase in the frequency of the CD56^dim^CD57^+^ NK cell subset was observed. These results can be explained by the fact that perforin and CD57 expression is likely upregulated across multiple NK cell subsets during acute DENV infection. Consequently, an increase in their expression can be predictive of infection without changing the distribution of the NK cell subsets.

Adult DENV infection led to increased expression of Fas across all myeloid cell subsets. This, along with the fact that adult NK cells upregulate Fas-L, suggests that adult NK cells may kill DENV-infected myeloid cells via the Fas-Fas-L pathway. CD14^+^CD16^−^ and CD141^+^ myeloid cells in DENV-infected children upregulated DR4/DR5 alone or in combination with Fas. Unfortunately, our NK cell CyTOF panel did not include the DR4/DR5 ligand, TNF-related apoptosis inducing ligand (TRAIL), and additional PBMC samples are not available. Therefore, we cannot determine whether this pathway is contributing to myeloid cell apoptosis in children. A previous study has shown an increase in the percentage of TRAIL^+^ NK cells as well as an increase in TRAIL expression during adult DENV infection (41), making it likely that TRAIL-mediated apoptosis is occurring.

Strikingly, acute DENV infection in adults led to fewer phenotypic changes in myeloid cells than in children. Of the markers whose expression increased during adult DENV infection, many would facilitate NK cell targeting. Besides Fas, two of the myeloid subsets expressed MICA/MICB and/or Nectin-2. MICA/MICB are ligands for the activating receptor NKG2D whose expression on NK cells was increased during adult DENV infection and was predictive of infection. Nectin-2 is a ligand for the activating receptor DNAM-1, which was expressed on the CD56^dim^perforin^bright^ NK cell subset. Many ligands for activating NK cell receptors were similarly upregulated during pediatric DENV infection. These included DR4/DR5, Fas, Nectin-2, as well as DNAM-1 ligand CD155 and NKG2D ligands ULBP-1,2,5,6. However, unlike what was observed in the adults, there was also significant induction of LLT-1, a ligand for the inhibitory receptor CD161, and HLA class I molecules, which are known to suppress the anti-DENV NK cell response (27–29). Only CD14^+^CD16^−^ adult myeloid cells demonstrated an increase in LLT-1 expression. Together, these findings suggest that during pediatric DENV infection, myeloid cells adopt a more NK cell-suppressive phenotype than adult myeloid cells. This potentially facilitates DENV-infected myeloid cell evasion of the NK cell response in children.

This study has limitations. The first is the small sample size of DENV-infected patients, particularly in our NK cell analysis. While the numbers may be modest, this data provides a solid foundation on which to design future studies comparing the adult and pediatric NK cell response to DENV infection. Another limitation was the lack of functional assessments, which did not allow us to directly evaluate whether adult NK cells are more responsive to DENV-infected cells than pediatric NK cells. Unfortunately, we do not have sufficient samples to perform such studies.

Overall, our results show that NK cells from pediatric and adult patients are uniquely impacted by acute DENV infection. We discovered that the frequency of total NK cells decreases in DENV-infected children, but not in DENV-infected adults. This decrease in pediatric NK cell frequency is accompanied by an increase in the abundance of a CD56^dim^perforin^bright^ NK cell subset. During adult DENV infection, NK cells develop an activated phenotype whereas the phenotype of pediatric NK cells is largely one of increased maturation. Similarly, adult myeloid cells upregulate ligands for activating NK cell receptors that may facilitate killing of DENV-infected cells by CD56^dim^perforin^bright^ NK cells and the Fas-Fas-L pathway. In contrast, myeloid cells from pediatric patients upregulate inhibitory ligands, which may suppress the NK cell response. These findings need to be confirmed with a larger DENV cohort that includes longitudinal samples. This will be critical to determining how the observed differences in the anti-DENV immune response in children and adults change across the different phases of infection. It would also be important to determine whether shifts in expression of specific markers or the frequencies of certain immune cell subsets correlate with disease severity. Such analyses may point to age-specific predictors of progression to severe DENV. Finally, functional studies comparing the ability of NK cells derived from children and adults to respond to DENV-infected cells are necessary to determine whether functional differences exist. Overall, this study provides insight into the differences between the NK cell response to pediatric and adult DENV infection which may have a bearing on disease severity.

## Acknowledgments

The authors would like to thank all health institutions, patients, their families, Luis Bonilla from the Blood Bank of Santo Tomas Hospital, Brechla Moreno, Yamilka Diaz, Mario Quijada, Yenesis Jordan, Arcelys Pitti, Zumara Chaverra, Jean-Paul Carrera, and all members of the Department of Virology and Biotechnology from ICGES. Thank you to the Stanford Human Immune Monitoring Center for use of their Helios machines as well as Lewis Lanier and Eva Harris for their helpful insights.

## Footnotes

### Author Contributions

Conceptualization, JM, AF, DB, LJS, CB, SL-V; Methodology, JM, AF, DB, LS, RV, DE, AA, OV, SH, SL-V; Formal Analysis, JM, AF, CB; Investigation, JM, DB, AF, CB, SL-V; Resources, DB, CB, SL-V; Data Curation, JM, AF, CB; Writing – Original Draft Preparation, JM, AF, CB; Supervision, SH, CB, SL-V; Project Administration, CB, SL-V; Funding Acquisition, DB, CB, SL-V.

---

**Supplementary Figure 1:** Live PBMC gating scheme.

**Supplementary Figure 2:** Undifferentiated febrile illness in pediatric patients uniquely affects the frequencies of specific immune cell subsets. **(A)** Results from a differential abundance test comparing the frequencies of immune cell subsets in children presenting with an undifferentiated febrile illness (green, n = 7) to healthy children (pink, n = 22). **(B)** Results from a differential abundance test comparing the frequencies of immune cell subsets in DENV-infected children (teal, n = 19) and children presenting with an undifferentiated febrile illness (green, n = 7). Subsets whose frequencies were significantly different (adjusted p-value < 0.05) between the two states are denoted by green boxes. The proportions of each cluster in each sample are represented by the normalized frequencies. Grey boxes correspond to under-representation and red boxes correspond to over-representation. The frequencies were scaled using arcsine-square-root transformation and then z-score normalized in each cluster **(A and B)**.

**Supplementary Figure 3:** NK cell gating scheme.

## References

1. 2009. EPIDEMIOLOGY, BURDEN OF DISEASE AND TRANSMISSION,. World Health Organization.

2. Hammond, S. N., A. Balmaseda, L. Pérez, Y. Tellez, S. I. Saborío, J. C. Mercado, E. Videa, Y. Rodriguez, M. A. Pérez, R. Cuadra, S. Solano, J. Rocha, W. Idiaquez, A. Gonzalez, and E. Harris. 2005. Differences in dengue severity in infants, children, and adults in a 3-year hospital-based study in Nicaragua. Am. J. Trop. Med. Hyg. 73: 1063–1070.

3. de Souza, L. J., L. Bastos Pessanha, L. Carvalho Mansur, L. Assed de Souza, M. Barbosa Tâmega Ribeiro, M. do Vale da Silveira, and J. T. Damian Souto Filho. 2013. Comparison of clinical and laboratory characteristics between children and adults with dengue. Braz. J. Infect. Dis. 17: 27–31.

4. Hanafusa, S., C. Chanyasanha, D. Sujirarat, I. Khuankhunsathid, A. Yaguchi, and T. Suzuki. 2008. Clinical features and differences between child and adult dengue infections in Rayong Province, southeast Thailand. Southeast Asian J. Trop. Med. Public Health 39: 252–259.

5. Namvongsa, V., C. Sirivichayakul, S. Songsithichok, P. Chanthavanich, W. Chokejindachai, and R. Sitcharungsi. 2013. Differences in clinical features between children and adults with dengue hemorrhagic fever/dengue shock syndrome. Southeast Asian J. Trop. Med. Public Health 44:772–779.

6. Winter, P. E., T. M. Yuill, S. Udomsakdi, D. Gould, S. Nantapanich, and P. K. Russell. 1968. An insular outbreak of dengue hemorrhagic fever. I. Epidemiologic observations. Am. J. Trop. Med. Hyg. 17: 590–599.

7. Chareonsook, O., H. M. Foy, A. Teeraratkul, and N. Silarug. 1999. Changing epidemiology of dengue hemorrhagic fever in Thailand. Epidemiol. Infect. 122: 161–166.

8. Kouri, G. P., M. G. Guzmán, J. R. Bravo, and C. Triana. 1989. Dengue haemorrhagic fever/dengue shock syndrome: lessons from the Cuban epidemic, 1981. Bull. World Health Organ. 67: 375–380.

9. Guzmán, M. G., G. Kouri, J. Bravo, L. Valdes, S. Vazquez, and S. B. Halstead. 2002. Effect of age on outcome of secondary dengue 2 infections. Int. J. Infect. Dis. 6: 118–124.

10. Díaz, Y., M. Chen-Germán, E. Quiroz, J.-P. Carrera, J. Cisneros, B. Moreno, L. Cerezo, A. O. Martinez-Torres, L. Moreno, I. Barahona de Mosca, B. Armién, R. Chen, N. Vasilakis, and S. López-Vergès. 2019. Molecular Epidemiology of Dengue in Panama: 25 Years of Circulation. Viruses 11.

11. Gamble, J., D. Bethell, N. P. Day, P. P. Loc, N. H. Phu, I. B. Gartside, J. F. Farrar, and N. J. White. 2000. Age-related changes in microvascular permeability: a significant factor in the susceptibility of children to shock? Clin. Sci. 98: 211–216.

12. Chau, T. N. B., N. T. Hieu, K. L. Anders, M. Wolbers, L. B. Lien, L. T. M. Hieu, T. T. Hien, N. T. Hung, J. Farrar, S. Whitehead, and C. P. Simmons. 2009. Dengue virus infections and maternal antibody decay in a prospective birth cohort study of Vietnamese infants. J. Infect. Dis. 200:1893–1900.

13. Kliks, S. C., S. Nimmanitya, A. Nisalak, and D. S. Burke. 1988. Evidence that maternal dengue antibodies are important in the development of dengue hemorrhagic fever in infants. Am. J. Trop. Med. Hyg. 38: 411–419.

14. Wichmann, O., S. Hongsiriwon, C. Bowonwatanuwong, K. Chotivanich, Y. Sukthana, and S. Pukrittayakamee. 2004. Risk factors and clinical features associated with severe dengue infection in adults and children during the 2001 epidemic in Chonburi, Thailand. Trop. Med. Int. Health 9: 1022–1029.

15. Simon, A. K., G. A. Hollander, and A. McMichael. 2015. Evolution of the immune system in humans from infancy to old age. Proc. Biol. Sci. 282: 20143085.

16. Prendergast, A. J., P. Klenerman, and P. J. R. Goulder. 2012. The impact of differential antiviral immunity in children and adults. Nat. Rev. Immunol. 12: 636–648.

17. Strauss-Albee, D. M., J. Fukuyama, E. C. Liang, Y. Yao, J. A. Jarrell, A. L. Drake, J. Kinuthia, R. R. Montgomery, G. John-Stewart, S. Holmes, and C. A. Blish. 2015. Human NK cell repertoire diversity reflects immune experience and correlates with viral susceptibility. Sci. Transl. Med. 7: 297ra115.

18. O’Sullivan, T. E., J. C. Sun, and L. L. Lanier. 2015. Natural Killer Cell Memory. Immunity 43: 634–645.

19. Green, S., S. Pichyangkul, D. W. Vaughn, S. Kalayanarooj, S. Nimmannitya, A. Nisalak, I. Kurane, A. L. Rothman, and F. A. Ennis. 1999. Early CD69 expression on peripheral blood lymphocytes from children with dengue hemorrhagic fever. J. Infect. Dis. 180: 1429–1435.

20. Azeredo, E. L. 2006. NK cells, displaying early activation, cytotoxicity and adhesion molecules, are associated with mild dengue disease. 345–356.

21. Keawvichit, R., L. Khowawisetsut, S. Lertjuthaporn, K. Tangnararatchakit, N. Apiwattanakul, S. Yoksan, A. Chuansumrit, K. Chokephaibulkit, A. A. Ansari, N. Onlamoon, and K. Pattanapanyasat. 2018. Differences in activation and tissue homing markers of natural killer cell subsets during acute dengue infection. Immunology 153: 455–465.

22. Petitdemange, C., N. Wauquier, H. Devilliers, H. Yssel, I. Mombo, M. Caron, D. Nkoghe, P. Debre, E. Leroy, and V. Vieillard. 2016. Longitudinal Analysis of Natural Killer Cells in Dengue Virus-Infected Patients in Comparison to Chikungunya and Chikungunya/Dengue Virus-Infected Patients. PLoS Negl. Trop. Dis. 10: 1–17.

23. Zimmer, C. L., M. Cornillet, C. Solà-Riera, K.-W. Cheung, M. A. Ivarsson, M. Q. Lim, N. Marquardt, Y.-S. Leo, D. C. Lye, J. Klingström, P. A. MacAry, H.-G. Ljunggren, L. Rivino, and N. K. Björkström. 2019. NK cells are activated and primed for skin-homing during acute dengue virus infection in humans. Nat. Commun. 10: 3897.

24. Hershkovitz, O., B. Rosental, L. A. Rosenberg, M. E. Navarro-Sanchez, S. Jivov, A. Zilka, O. Gershoni-Yahalom, E. Brient-Litzler, H. Bedouelle, J. W. Ho, K. S. Campbell, B. Rager-Zisman, P. Despres, and A. Porgador. 2009. NKp44 receptor mediates interaction of the envelope glycoproteins from the West Nile and dengue viruses with NK cells. J. Immunol. 183: 2610–2621.

25. Naiyer, M. M., S. A. Cassidy, A. Magri, V. Cowton, K. Chen, S. Mansour, H. Kranidioti, B. Mbirbindi, P. Rettman, S. Harris, L. J. Fanning, A. Mulder, F. H. J. Claas, A. D. Davidson, A. H. Patel, M. A. Purbhoo, and S. I. Khakoo. 2017. KIR2DS2 recognizes conserved peptides derived from viral helicases in the context of HLA-C. 2: 1–12.

26. Costa, V. V., W. Ye, Q. Chen, M. M. Teixeira, P. Preiser, E. E. Ooi, and J. Chen. 2017. Dengue Virus-Infected Dendritic Cells, but Not Monocytes, Activate Natural Killer Cells through a Contact-Dependent Mechanism Involving Adhesion Molecules. MBio 8.

27. Glasner, A., E. Oiknine-Djian, Y. Weisblum, M. Diab, A. Panet, D. G. Wolf, and O. Mandelboim. 2017. Zika Virus Escapes NK Cell Detection by Upregulating Major Histocompatibility Complex Class I Molecules. J. Virol. 91.

28. Hershkovitz, O., A. Zilka, A. Bar-Ilan, S. Abutbul, A. Davidson, M. Mazzon, B. M. Kummerer, A. Monsoengo, M. Jacobs, and A. Porgador. 2008. Dengue Virus Replicon Expressing the Nonstructural Proteins Suffices To Enhance Membrane Expression of HLA Class I and Inhibit Lysis by Human NK Cells. J. Virol. 82: 7666–7676.

29. McKechnie, J. L., D. Beltrán, A. Pitti, L. Saenz, A. B. Araúz, R. Vergara, E. Harris, L. L. Lanier, C. A. Blish, and S. López-Vergès. 2019. HLA Upregulation During Dengue Virus Infection Suppresses the Natural Killer Cell Response. Front. Cell. Infect. Microbiol. 9: 268.

30. Zhao, Y., M. Amodio, B. Vander Wyk, B. Gerritsen, M. M. Kumar, D. van Dijk, K. Moon, X. Wang, A. Malawista, M. M. Richards, M. E. Cahill, A. Desai, J. Sivadasan, M. M. Venkataswamy, V. Ravi, E. Fikrig, P. Kumar, S. H. Kleinstein, S. Krishnaswamy, and R. R. Montgomery. 2020. Single cell immune profiling of dengue virus patients reveals intact immune responses to Zika virus with enrichment of innate immune signatures. PLoS Negl. Trop. Dis. 14: e0008112.

31. Durbin, A. P., M. J. Vargas, K. Wanionek, S. N. Hammond, A. Gordon, C. Rocha, A. Balmaseda, and E. Harris. 2008. Phenotyping of peripheral blood mononuclear cells during acute dengue illness demonstrates infection and increased activation of monocytes in severe cases compared to classic dengue fever. Virology 376: 429–435.

32. Patumanond, J., C. Tawichasri, and S. Nopparat. 2003. Dengue hemorrhagic fever, Uttaradit, Thailand. Emerg. Infect. Dis. 9: 1348–1350.

33. Guzman, M. G., and G. Kouri. 2003. Dengue and dengue hemorrhagic fever in the Americas: lessons and challenges. J. Clin. Virol. 27: 1–13.

34. Messina, J. P., O. J. Brady, N. Golding, M. U. G. Kraemer, G. R. W. Wint, S. E. Ray, D. M. Pigott, F. M. Shearer, K. Johnson, L. Earl, L. B. Marczak, S. Shirude, N. Davis Weaver, M. Gilbert, R. Velayudhan, P. Jones, T. Jaenisch, T. W. Scott, R. C. Reiner Jr, and S. I. Hay. 2019. The current and future global distribution and population at risk of dengue. Nat Microbiol 4: 1508–1515.

35. Chandele, A., J. Sewatanon, S. Gunisetty, M. Singla, N. Onlamoon, R. S. Akondy, H. T. Kissick, K. Nayak, E. S. Reddy, H. Kalam, D. Kumar, A. Verma, H. Panda, S. Wang, N. Angkasekwinai, K. Pattanapanyasat, K. Chokephaibulkit, G. R. Medigeshi, R. Lodha, S. Kabra, R. Ahmed, and K. Murali-Krishna. 2016. Characterization of Human CD8 T Cell Responses in Dengue Virus-Infected Patients from India. J. Virol. 90: 11259–11278.

36. Naranjo-Gomez, J. S., J. A. Castillo, P. A. Velilla, M. Rojas, and D. Castano. 2015. Frequency and phenotype alterations in monocyte subsets from dengue patients. In Front. Immunol. Conference Abstract: IMMUNOCOLOMBIA2015-11th Congress of the Latin American Association of Immunology-10o. Congreso de la Asociación Colombiana de Alergia, Asma e Inmunología. doi: 10.3389/conf.fimmu vol. 180.

37. Kwissa, M., H. I. Nakaya, N. Onlamoon, J. Wrammert, F. Villinger, G. C. Perng, S. Yoksan, K. Pattanapanyasat, K. Chokephaibulkit, R. Ahmed, and B. Pulendran. 2014. Dengue Virus Infection Induces Expansion of a CD14 CD16 Monocyte Population that Stimulates Plasmablast Differentiation. Cell Host & Microbe 16: 115–127.

38. Singla, M., M. Kar, T. Sethi, S. K. Kabra, R. Lodha, A. Chandele, and G. R. Medigeshi. 2016. Immune Response to Dengue Virus Infection in Pediatric Patients in New Delhi, India—Association of Viremia, Inflammatory Mediators and Monocytes with Disease Severity. PLoS Negl. Trop. Dis. 10: 1–25.

39. Boyette, L. B., C. Macedo, K. Hadi, B. D. Elinoff, J. T. Walters, B. Ramaswami, G. Chalasani, J. M. Taboas, F. G. Lakkis, and D. M. Metes. 2017. Phenotype, function, and differentiation potential of human monocyte subsets. PLoS One 12: e0176460.

40. Shao, J.-Y., W.-W. Yin, Q.-F. Zhang, Q. Liu, M.-L. Peng, H.-D. Hu, P. Hu, H. Ren, and D.-Z. Zhang. 2016. Siglec-7 defines a highly functional natural killer cell subset and inhibits cell-mediated activities. Scand. J. Immunol. 84: 182–190.

41. Gandini, M., F. Petitinga-Paiva, C. F. Marinho, G. Correa, L. M. De Oliveira-Pinto, L. J. D. Souza, R. V. Cunha, C. F. Kubelka, and E. L. D. Azeredo. 2017. Dengue Virus Induces NK Cell Activation through TRAIL Expression during Infection. Mediators Inflamm. 2017.

42. Lopez-Vergès, S., J. M. Milush, S. Pandey, V. A. York, J. Arakawa-Hoyt, H. Pircher, P. J. Norris, D. F. Nixon, and L. L. Lanier. 2010. CD57 defines a functionally distinct population of mature NK cells in the human CD56dimCD16+ NK-cell subset. Blood 116: 3865–3874.

43. Le Garff-Tavernier, M., V. Béziat, J. Decocq, V. Siguret, F. Gandjbakhch, E. Pautas, P. Debré, H. Merle-Beral, and V. Vieillard. 2010. Human NK cells display major phenotypic and functional changes over the life span. Aging Cell 9: 527–535.

44. Nielsen, C. M., M. J. White, M. R. Goodier, and E. M. Riley. 2013. Functional Significance of CD57 Expression on Human NK Cells and Relevance to Disease. Front. Immunol. 4: 422.

45. Bunn, A., and M. Korpela. 2019. An introduction to dplR..

46. McInnes, L., J. Healy, and J. Melville. 2018. UMAP: Uniform Manifold Approximation and Projection for Dimension Reduction. arXiv [stat.ML].

47. Van Gassen, S., B. Callebaut, M. J. Van Helden, B. N. Lambrecht, P. Demeester, T. Dhaene, and Y. Saeys. 2015. FlowSOM: Using self-organizing maps for visualization and interpretation of cytometry data. Cytometry A 87: 636–645.

48. Wilkerson, M. D., and D. N. Hayes. 2010. ConsensusClusterPlus: a class discovery tool with confidence assessments and item tracking. Bioinformatics 26: 1572–1573.

49. Melville, J. The Uniform Manifold Approximation and Projection (UMAP) Method for Dimensionality Reduction [R package uwot version 0.1.8]..

50. Weber, L. M., M. Nowicka, C. Soneson, and M. D. Robinson. 2019. diffcyt: Differential discovery in high-dimensional cytometry via high-resolution clustering. Commun Biol 2: 183.

51. Seiler, C., L. M. Kronstad, L. J. Simpson, M. Le Gars, E. Vendrame, C. A. Blish, and S. Holmes. 2019. Uncertainty Quantification in Multivariate Mixed Models for Mass Cytometry Data. arXiv [stat.AP].

52. Kronstad, L. M., C. Seiler, R. Vergara, S. P. Holmes, and C. A. Blish. 2018. Differential Induction of IFN-α and Modulation of CD112 and CD54 Expression Govern the Magnitude of NK Cell IFN-γ Response to Influenza A Viruses. J. Immunol. 201: 2117–2131.

53. Efron, B., and R. Tibshirani. 1993. An Introduction to the Bootstrap. New York: Chapman & Hall/CRC.

