## Supplementary Table I for "Mass cytometry analysis of the NK cell receptor-ligand repertoire reveals unique differences between dengue-infected children and adults"

| **Specificity** | **Clone** | **Isotope** |
| --- | --- | --- |
| HLA-DR | L243 | 89Y |
| CD45 | HI30 | 102Pd |
| CD45 | HI30 | 104Pd |
| CD45 | HI30 | 106Pd |
| CD45 | HI30 | 108Pd |
| CD19 | SJ25-C1 | Qdot® 655 (112Cd-114Cd) |
| CD3 | UCHT1 | 115In |
| CD20 | 2H7 | 141Pr |
| CD163 | GH1/61 | 142Nd |
| Pan HLA class I | W6/32 | 143Nd |
| CD7 | CD7-6B7 | 144Nd |
| CD8 | SK1 | 145Nd |
| CD48 | BJ40 | 146Nd |
| BDCA-2 (CD303) | 201A | 147Sm |
| ICAM-1 | HA58 | 148Nd |
| LLT-1 | 402659 | 149Sm |
| Flavi E protein | D1-4G2-4-15 | 150Nd |
| CD4 | OKT4 | 151Eu |
| CD64 | 10.1 | 152Sm |
| HLA-B/C | DT9 | 153Eu |
| CCR2 | K036C2 | 154Sm |
| HLA-E | 3D12 | 155Gd |
| Fas (CD95) | DX2 | 156Gd |
| Nectin-1 | R1.302 | 157Gd |
| MICA/B | 159227/236511 | 158Gd |
| DR4/5 | DJR1/DJR2-2 | 159Tb |
| CD1c | L161 | 160Gd |
| ULBP-1,2,5,6 | 170818/165903 | 161Dy |
| CD11c | Bu15 | 162Dy |
| NS3 | E1D8 | 163Dy |
| Nectin-2 | TX31 | 164Dy |
| CD155 | SKII.4 | 165Ho |
| HLA-Bw4 | REA274 | 166Er |
| CD32 | IV.3 | 167Er |
| HLA-Bw6 | REA143 | 168Er |
| CD14 | M5E2 | 169Tm |
| CD11b | ICRF44 | 170Er |
| LFA-3 | TS2/9 | 171Yb |
| CD33 | WM53 | 172Yb |
| CD141 (BDCA-3) | 1A4 | 173Yb |
| CD56 | NCAM16.2 | 174Yb |
| CD86 | IT2.2 | 175Lu |
| B7-H6 | 875001 | 176Yb |
| DNA-1/DNA-2 | NA | 191Ir/193Ir |
| Cisplatin | NA | 194Pt/195Pt |
| CD16 | 3G8 | 209Bi |

**Supplementary Table I:** PBMC CyTOF panel.
