## Supplementary figures and images for "Mass cytometry analysis of the NK cell receptor-ligand repertoire reveals unique differences between dengue-infected children and adults"

### Supplementary Figure 1

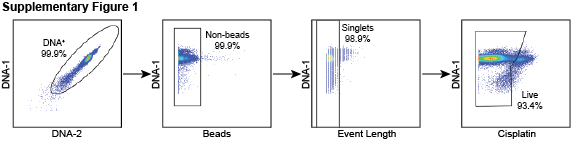

### Supplementary Figure 2

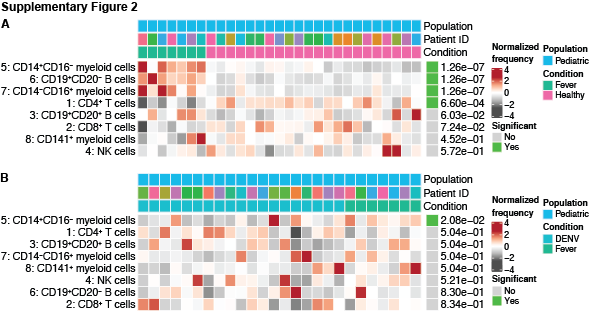

### Supplementary Figure 3

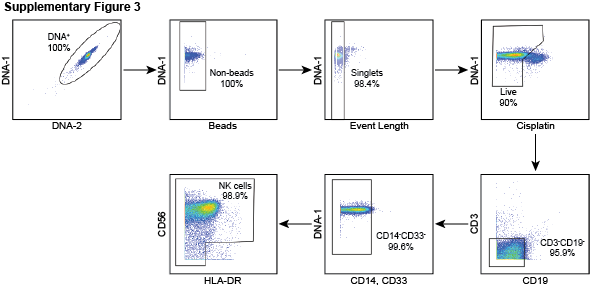
