## Supplementary Table II for "Mass cytometry analysis of the NK cell receptor-ligand repertoire reveals unique differences between dengue-infected children and adults"

| **Specificity** | **Clone** | **Isotope** |
| --- | --- | --- |
| CD57 | HCD57 | 89Y |
| HLA-DR | Tu36 | Qdot® 655 (112Cd-114Cd) |
| CD3 | UCHT | 115In |
| CD38 | HIT2 | 141Pr |
| CD69 | FN50 | 142Nd |
| CD33 | WM53 | 143Nd |
| CD14 | M5E5 | 143Nd |
| CD2 | RPA-2.10 | 144Nd |
| CD19 | HIB19 | 146Nd |
| CD8 | SK1 | 147Sm |
| FcRγ | Polyclonal | 148Nd |
| CD4 | SK3 | 149Sm |
| Syk | 4D10.2 | 150Nd |
| CD62L | DREG-56 | 151Eu |
| Ki-67 | Ki-67 | 152Sm |
| KIR2DS4 | 179315 | 153Eu |
| KIR2DS2 | Polyclonal | 154Sm |
| NKp46 | 9E2 | 155Gd |
| NKG2D | 1D11 | 156Gd |
| TIGIT | 741182 | 157Gd |
| 2B4 | C1.7 | 158Gd |
| DNAM-1 | DX11 | 159Tb |
| FAS-L | NOK-1 | 160Gd |
| NKp30 | P30-15 | 161Dy |
| Siglec-7 | S7.7 | 162Dy |
| NKG2C | 134522 | 163Dy |
| NKp44 | P44-8 | 164Dy |
| TACTILE (CD96) | NK92.39 | 165Ho |
| KIR2DL1 | 143211 | 166Er |
| CD94 | DX22 | 167Er |
| CXCR6 | K041E5 | 168Er |
| PD-1 | EH12.2H7 | 169Tm |
| KIR2DL5 | UP-R1 | 170Er |
| NKG2A | 131411 | 171Yb |
| NTB-A | NT-7 | 172Yb |
| KIR3DL1 | DX-9 | 173Yb |
| CD56 | NCAM16.2 | 174Yb |
| KIR2DL3 | 180701 | 175Lu |
| Perforin | B-D48 | 176Yb |
| CD16 | 3G8 | 209Bi |

**Supplementary Table II:** NK cell CyTOF panel.
